# Pharmacovigilance or patient safety: analysis from a patient safety program

**DOI:** 10.1101/399923

**Authors:** Yira Constanza Cortázar C, José Gilberto Orozco D, José Julián López G

## Abstract

**Purpose:** drugs are the common point of pharmacovigilance and patient safety programs. Despite using a common language, the same epidemiological method and legislation that requires the operation of the two programs, there does not seem to be a clear relationship between them.

**Methodology:** observational descriptive cross sectional study of the reports database from an institutional patient safety program. Medication errors were classified according to the document *The Conceptual Framework for the International Classification for Patient Safety* (ICPS) WHO 2009. Adverse Reactions (ADR) were classified according to Uppsala Monitoring Center.

**Results:** the omission of drugs or doses was the most frequent error with 42.8% followed by ADRs (20.9%). No harm incidents corresponded to 61.2% and the remaining 38.8% was represented in near missincidents and no harm incidents. There were included 41 ADR and 15 therapeutic failures corresponding to a point-prevalence of 57 ADR/10,000 patients-year and 28.6% (56/196) of reports related to drugs. Phlebitis is the most frequently reported with 23, 7% followed by hypersensitivity reactions with 18.4% and excessive neuromuscular blockade with 13.1%.

**Conclusions:** considering time, level of care and number of bed, ADR prevalence seem low. A very important proportion of reports corresponding to near miss incidents or no harm incidents is not taken into account by the security managers, losing a valuable risk management opportunity in the patient safety programs.

## Introduction

Although by 1848 an attempt had already been made to report adverse drug reaction (ADR) suspicions because of a young woman died by the administration of chloroform during surgery in England (1), it was only up to 1960 when the first pharmacovigilance (PV) systems were originated as a result of the outbreak of phocomelia caused by the administration of thalidomide (2). After this, a series of articles have been published showing the harm caused by drugs: Talley identified in 1974 that 2,9% of admissions to the medical service were for this cause and 6, 2% of these patients died (3). Subsequently, Manasse stated that by 1987 drug mortality affected 12,000 Americans and morbidity reached 15,000 hospitalizations. He also coined the term *Drug misadventuring* to describe negative drug experiences that he considered a public policy problem derived from the excessive use of drugs and error-prone preparation and distribution systems (4, 5). Lazarou affirm in 1998 that ADR were between the fourth and sixth leading cause of death (6).

According to the current definition of the World Health Organization (WHO), PV is “the science and activities relating to the detection, assessment, understanding and prevention of adverse effects or any other possible drug-related problems…”. (7). By 2012, the national pharmacovigilance program in Colombia considered that PV should study the problems related with the use of drugs and its effects in society for preventing and resolving them (8, 9). Today, regulatory agencies work to balance access to drugs with safety concerns, in line with their mission to protect the public health. However, in the first years of the 21st century, the safety of prescribed drugs caught the attention of the public because of some drugs withdrawal, delays in warning the public about important risks and the approval of drugs without enough attention to safety. (10)

Not only ADRs are part of daily concerns of health professionals, there is also interest in others risks of health care such as nosocomial infections, complications of the clinical course and medication errors (ME). There have been numerous studies where it was estimated that the incidence rate of adverse events in Spanish hospitals were 9.3%, 46.6 were considered preventable; 37.4% were related with medication which 34.8% were preventable. (11).

The document *To err is human* identified that about 98,000 people die in a year because of errors in health care occurring in hospitals. After this document, the WHO in 2004 pointed out to the governments of the world the need to establish programs that guarantee safe actions in the patient care and suggests a global strategy to fulfill this purpose (12, 13). Colombia has not been unaware of this situation and through the Ministry of Social Protection urges institutions to implement and continuously validate a Patient Safety Program (PSP), which guarantees the best conditions for the healthcare of Colombian people and for which it has issued specific regulations (14, 15).

PSPs in their philosophy extend the approach of safety in patient care and include in their objectives the inspection of activities related to healthcare, such as skin integrity, prevention of falls, control of medical devices, and surveillance of blood derivatives, among others. These programs incorporate in their concept the inherent risk of health care service and part of their analysis includes the evaluation of the causes of errors or failures in the system that will allow establishing corrective actions in future risk situations for the patient. IBEAS study identified that a 10.5% of the patients presented an adverse event and at least half of these adverse events to drugs could be prevent (16).

Despite these two programs led by the WHO, the negative consequences for the patients’ health are still far from being controlled or minimized due, among other reasons, to the current biomedical model that aims to solve health issues with medical interventions, in which drugs are an essential part, and a neoliberal economy that turned health into a business model (17, 18).

One of the common points of PVP and PSP are drugs. Despite the use of common language such as ADR, ME and the same epidemiological method (risk approach) added to the fact that in Colombian legislation it is required to have the two programs in order to achieve the certification of institutions for provision of health services to the population, there seems to be no clear articulation between them and it has even been identified that there is a significant variability in PVP, which limits the comparability between different information systems(19). From an institutional PSP, this study characterized the drug related reports sent to the program, identified the differences of their classification.

## Materials and methods

An observational descriptive cross sectional study was carried out with a retrospective collection of information, reports of the institutional PSP application were included during 2015, and there were excluded duplicate reports, invalid (report presenting any inconsistency), tests (information is introduced to verify the integrity of the application), related to the infrastructure of the hospital, and personal complaints. The reports were classified according to the categories established in the document *The Conceptual Framework for the International Classification for Patient Safety* (ICPS) of WHO 2009 (20). For the ADR classification, the tools suggested by the Uppsala Monitoring Center were used, such system/organ affected and seriousness (21).

The ethics committee approved the study conforms to the Declaration of Helsinki and resolution 008430 of 1993 according to the specified in the article 11 of chapter I. The present study is a risk-free investigation therefore written informed consent is not required.

## Results

According to the database used in the study, 1481 safety cases were reported in 2015. After the preliminary analysis, 439 (29.6%) reports were discarded, because they fulfilled some excluded criteria, leaving 1042 total valid reports of which 196 (18.8%) were identified and are related to drugs and make up the common nucleus between PSP and PVP. Fig 1 shows the process and the results of the initial evaluation.

**Figure 1.**
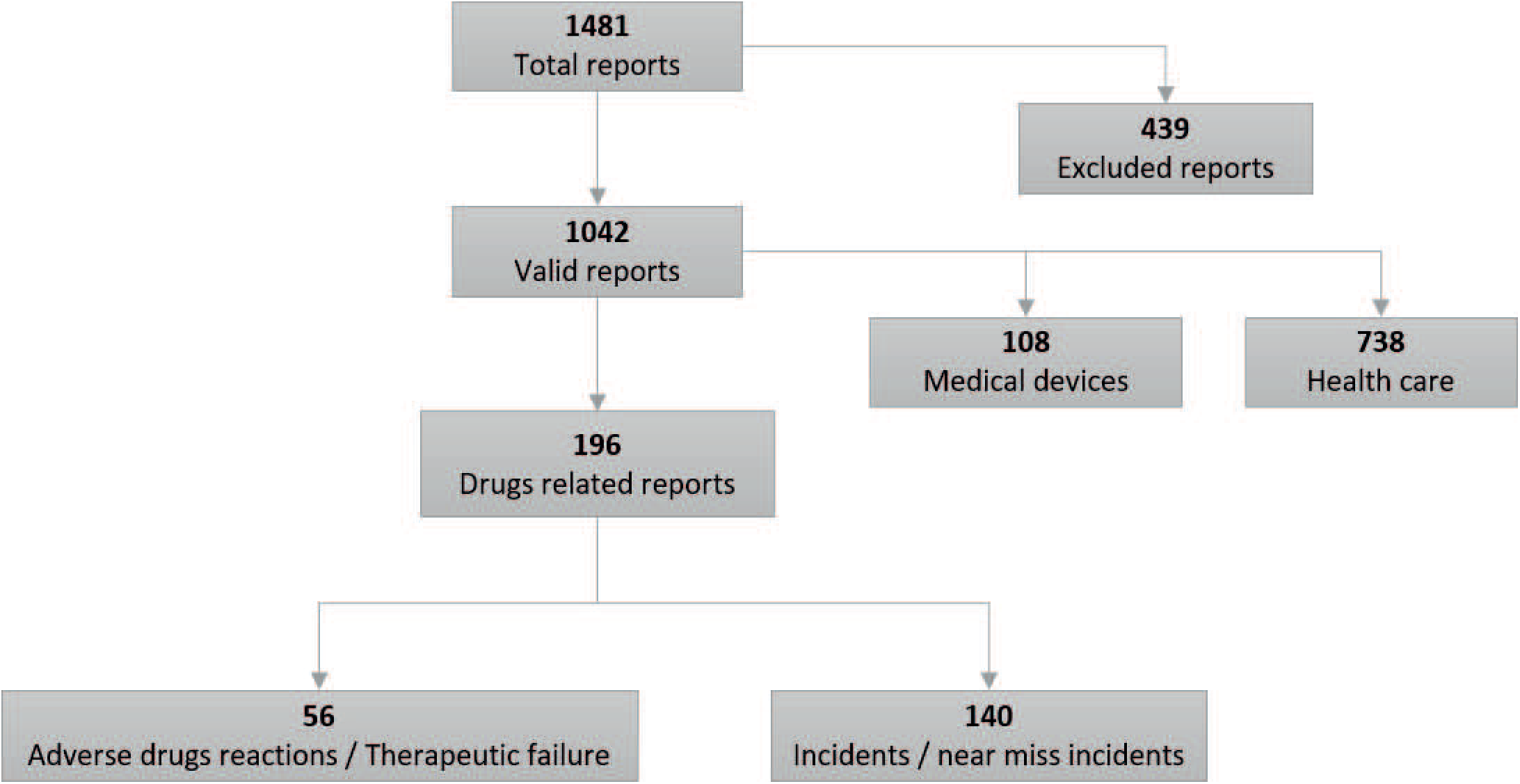
Flow diagram of the selection and classification of the reports. Steps for the selection of the reports included in the present study. Only those related to medications were included.

### PSP: medication errors

Table 1 shows the distribution of reports according to ICPS, where it can be observed that 42.8% (84/196) correspond to omission of drugs or dose (category 9), followed by 20.9% (41/196) related to ADR (category 11). Sixteen reports could not be classified in any category so it was necessary to create two additional categories: “therapeutic failure” (TF) (category 12) in which 15 reports were identified, and “unclassifiable” (category 20) corresponding to an event of wrong adjustment.

**Table 1.** Distribution of errors according to ICPS

| Error | Category | N | % |
| --- | --- | --- | --- |
| Wrong drug | 1 | 14 | 7,14 |
| Wrong patient | 2 | 3 | 1,53 |
| Wrong dose/strength of frequency | 3 | 10 | 5,10 |
| Wrong formulation or presentation | 4 | 4 | 2,04 |
| Wrong route or quantity | 5 | 1 | 0,51 |
| Wrong Dispensing Label/Instruction | 6 | 8 | 4,08 |
| Contraindication | 7 | 2 | 1,02 |
| Wrong storage | 8 | 13 | 6,63 |
| Omitted medicine or dose | 9 | 84 | 42,86 |
| Expired medicine | 10 | 0 | 0 |
| Adverse drug reaction | 11 | 41 | 20,92 |
| Therapeutic failure | 12 | 15 | 7,65 |
| Not classifiable | 20 | 1 | 0,51 |
|  | Total | 196 | 100 |

Classification of medication errors according to ICPS. The last two classifications (12 and 20) were created for the present study.

Reports were constituted by 61.2% (120/196) of harmful incidents and the remaining 38.8% (76/196) of the reports were represented near miss incidents and no harm incidents. Harmful incidents were distributed in ADRs (41/120), TF (15/120) and harm related to surgery delay, additional tests, drug rupture or harm or strict monitoring (64/120) are included. Of these events, 68% (82/120) corresponded to moderate harm.

### PVP: Adverse drug reactions

There were identified 41 ADRs and 15 TF that correspond to 5.4% (56/1040) of total reports and to 28.6% (56/196) of reports related to drugs. Table 2 shows that the most affected organ-system was the skin with a 42.8% (24/56), followed by the central nervous system with a 23.2% (13/56)

**Table 2.** Distribution of ADR according to WHO

| Drug | ADR | Classification | Affected System | N |
| --- | --- | --- | --- | --- |
| Dexmedetomidine | Agitation, anxiety | Moderate | Nervous | 1 |
| Rocuronium | Neuromuscular block | Serious | Musculoskeletal | 5 |
| Oxygen | hypertensive crisis | Serious | Cardiovascular | 1 |
| Noradrenaline | Emesis | Moderate | Gastrointestinal | 1 |
| No medication related | Emesis | Moderate | Gastrointestinal | 2 |
| Haloperidol | Extrapyramidalism | Moderate | Nervous | 1 |
| No medication related | Phlebitis | Moderate | Skin and its appendages | 11 |
| Polymyxin B | Phlebitis | Moderate | Skin and its appendages | 1 |
| Vancomycin | Phlebitis | Moderate | Skin and its appendages | 2 |
| Phenytoin | Phlebitis | Moderate | Skin and its appendages | 1 |
| Insulin glargine | hypoglycemia | Moderate | Endocrine | 1 |
| Prazosin | Hypotension | Moderate | Cardiovascular | 1 |
| Certolizumab | Reverse psoriasis | Moderate | Skin and its appendages | 1 |
| Cefazolin | Skin rash | Moderate | Skin and its appendages | 1 |
| Dipyrrone | Hypersensitivity reaction | Moderate | Skin and its appendages | 1 |
| Phenytoin | Hypersensitivity reaction | Moderate | Skin and its appendages | 1 |
| Hyoscine | Hypersensitivity reaction | Moderate | Skin and its appendages | 1 |
| Levetiracetam | Hypersensitivity reaction | Moderate | Skin and its appendages | 1 |
| Morphine | Hypersensitivity reaction | Moderate | Skin and its appendages | 1 |
| Piperacillin<br>tazobactam | Hypersensitivity reaction | Moderate | Skin and its appendages | 1 |
| Rituximab | Hypersensitivity reaction | Moderate | Skin and its appendages | 1 |
| Heparin | Overanticoagulation | Serious | Blood | 1 |
| Warfarin | Overanticoagulation | Serious | Blood | 3 |
| Midazolam | Therapeutic failure | Moderate | Nervous | 6 |
| Dexmedetomidine | Therapeutic failure | Moderate | Nervous | 1 |
| Fentanyl | Therapeutic failure | Moderate | Nervous | 4 |
| Labetalol | Therapeutic failure | Moderate | Cardiovascular | 4 |
| Total |  |  |  | 56 |

Classification of the ADR according to the Uppsala Monitoring Center, taking into account the severity and the affected organ-system

During the study period, 9900 patients were attended, which shows an incidence of ADR of 0.56% (456/9900). According to the organ-system affected, the skin and appendages disorders occupy the first place with 42.8% (24/56) followed by the nervous system with 23.2% (13/56). According to the seriousness of ADR, most of them were moderate with 82.2% (46/56) and the remaining 17.8% (10/56) were serious.

In the results of the causality analysis with the Naranjo algorithm (68), a score of 7 was obtained in all cases, with the exception of rocuronium, which was assigned an additional point (+8) because of the fact that sugammadex was used to reverse the blockade, however, they all go into the PROBABLE category.

### Drugs involved in the reports

The drugs classification by anatomical group was carried out according to the Anatomical Therapeutical Chemic (ATC) classification (22), in which the group of drugs most frequently involved in the reports were those of the N group with 30.4% (17/56) followed by the group M and C each with 10.7% (6/56). Other groups with low percentage were L and A. It was necessary to create “category X” (without specific drug), which corresponds to the report of an event where a problem is mentioned without specifically describing the international common denomination.

## Discussion

The present study identified that 1 out of 4 reports were rejected, this can be explained by some of the following reasons: incomplete socialization of the program, high healthcare burden, high professionals and students turnover (university hospital). These invalid reports have a negative impact on the program, because the classification of the reports is done manually and increases the reading time.

Another finding was a higher frequency of incidents reports related with care (70.8%), followed by those related to drugs (18.8%) and, finally, with medical devices (10.4%). Given that the interest of this work corresponds to drugs, it can be mentioned that the frequency of reports related to this supply is within the numbers identified in studies such as ENEAS (adverse events in hospitals in Spain, with 37.4%) (11), IBEAS (adverse events in hospitals in Latin America, with 8.23%) (16) and SYREC (adverse events in Spanish intensive care units, with 24.6%) (23).

Regarding the reports classification, it can be affirmed that about half correspond to category 9 of the ICPS (drugs omission) and another important percentage is related to categories 1 and 7 (inadequate conservation conditions and wrong medication). These findings may be a reflection of the lack of a drug distribution system, whose purpose is, precisely, to reduce these errors (24).

A study conducted in the hospitalization service of a clinic in Cali (Colombia) identified that in a period of 20 days the drugs omission was the most common error, although it does not mention which classification was used (25). Another study conducted by Machado et al. during 8 years in ambulatory pharmacies in different cities of Colombia identified that most of the errors were related to dispensation (26). However, it is necessary to emphasize that the comparison of these results with other studies is complicated by the use of different types of classification and different settings of study (community, hospital, ambulatory, etc.). For example, a recent report in England identifies the potentially inappropriate prescription as an error, as well as administration, monitoring and dispensing errors (27).

According to the typology of the reportable events, it is identified that more than half of the reports correspond to harmful incidents. Regarding the harmful incidents, it is necessary to clarify that from the perspective of PSP, these harms are not always related to the affectation of human biology (which in PV would be called ADR), but also include another type of harm that affects other areas of the person or the health system (surgery delay, intensive monitoring, pharmaceutical product damage, etc.). The development and consolidation of the program may lead to the predominance of the near miss incidents report, since they are closer to prevention and constitute the first sign of incidents that may or not lead to harm. This high proportion of harmful incidents reports may be related to the fact that people tend to report those events that cause harm, since those that do not bring consequences can be considered “normal”, not worthy of notification or a combination of the “seven capital sins of underreporting”, which will be discussed later (28).

According to this consideration, it draws attention that a little less than half of these incidents correspond to ADR and therefore were included in the activities of PVP, while the remaining percentage of incidents that also caused harm were not considered. Additionally, reports of incidents that did not cause harm (no harm incidents, near miss incidents) are not analyzed, mainly due to high workload, which means that an important opportunity to manage the risk is lost, since it should not wait for it to appear ADR to inform and then analyze. This is one of the most relevant findings of the present study, since it demonstrates that PV should also deal with errors or infractions in order to be corrected and prevented. Some authors have identified this need and make a call so that ME are taken into account in PVP (29, 30). It is therefore necessary to visualize these findings as an opportunity for improvement, first identifying that the scope of the two programs pose as a challenge the non-duplicity of efforts, and, as an opportunity, not leaving problems unattended, specifically the near miss incidents and no harm incidents.

In the referenced studies, it was not possible to identify the results in terms of no harm incidents or near miss-incidents. Some of them describe the results in terms of ADR or ME without taking into account the infractions and other reportable events mentioned at the beginning of the paragraph. It is not possible to make a direct comparison with other studies related to the subject for the following reasons: equal or very similar terms to refer different things (adverse event in PV vs. adverse event in PS); different terms to refer the same (adverse event or preventable ADR in PV vs. ME in PS); lack of knowledge by PV programs that only look at ADR and, finally, some discrepancies in the harmonization of terms used in the Colombian regulation vs. WHO (incident related to patient safety according to WHO 2009 and incident according to Minprotection 2008 and 2009).

In relation to the PVP and according to the institutional application, a 0.56% prevalence of ADR was estimated during a year, a number that seems low for a fourth-level hospital with 223 hospital beds and 9900 inpatients in the different services during a year, if it is considered that the studies indicate that in this type of hospitals it is presented between 10 to 20% of ADR, of which between a 10 to 20% are classified serious and the 0.5 to 0.9%% are mortal (11, 16). The results of the present study may also be related to underreporting, one of the main problems of passive pharmacovigilance (31). A document prepared by Varallos et al. explains that this, under report, occurs for what has been called “the seven capital sins of underreporting”: 1) consider that serious ADR are well documented, 2) fear of being involved in legal proceedings, 3) guilty feeling for have been responsible for the harm to the patient, 4) ambition of a group for publish serious cases, 5) lack of knowledge about how to make the notification, 6) insecurity about the report of ADR and, 7) indifference, lack of interest, time or another excuse to postpone notification. The main causes of ADR underreporting found in the studies included in a systematic review were ignorance and insecurity, findings related to the low knowledge of professionals on the activities of drugs safety analysis; the authors propose that professional notifications can be promoted through educational interventions aimed to clarify their importance (28). For the present study, this could also be the cause of poor registration in the institutional application and invalid reports.

The drugs involved in the reports differ from the results found by Machado et al. in Colombia (32–33), De las Salas et al. in two pediatric hospitals for 6 months (34), Moscoso et al. in a second-level hospital in Bogotá for 3 months (35) and a study by Chaves in 31 second-level institutions in the city of Bogotá for a year (36), where antibiotics are among the drugs that report the most ADR, although they agree that the skin it is the most affected organ-system. Although some similarities are found in the afore mentioned, differences are also found in some of the results, this can be explained by the type of institution and the methodology used for the identification (passive vs. active).

Despite the wide dissemination and published studies related to patient safety, no similar studies were found in the bibliographic review carried out for this study that attempted to reflect on the articulation of PV and PS programs in specific hospitals. However, documents such as the one written by the WHO in 2014 and the EMA in 2015 allow us to deduce that studying the real articulation of these programs is at the heart of the PV and PS programs (29, 37). For almost 20 years, several authors have discussed the need for a change in the scope, approach or methods used to perform PV. It is possible that these findings are the result of the movement on patient safety or of society’s need to counteract the growing outbreak of drug-induced iatrogenic (38–44).

There is a need to broaden the vision of health surveillance systems to include aspects such as drug-related problems, ME and ADR. It has not yet been possible to integrate and incorporate these terms into a single program, perhaps for reasons that range from the purely philosophical, to a predominance of positivism, passing through political and economic causes whose analysis goes beyond the scope of this research (45–47).

## Acknowledgments

The authors thank Universidad Nacional de Colombia

## References

1. Routledge P. 150 years of pharmacovigilance. Lancet. 1998; 351(9110):1200–1.

2. Strom BL. *What is Pharmacoepidemiology?* In: Brian L. Strom SEK, Sean Hennessy, editor. Pharmacoepidemiology. 5th ed. Chichester, UK: John Wiley & Sons, Ltd; 2012.

3. Talley RB, Laventurier MF. Drug-induced illness. JAMA. 1974;229(8):1043-.

4. Manasse H. Medication use in an imperfect world: Drug misadventuring as an issue of public policy part 1. Am J Hosp Phar. 1989;46(1):929–44.

5. Manasse H. Medication use in an imperfect world: Drug misadventuring as an issue of public policy part 2. AM J Hosp Pharm. 1989;46(1):1141–52.

6. Lazarou J, Pomeranz BH, Corey PN. Incidence of adverse drug reactions in hospitalized patients: a meta-analysis of prospective studies. JAMA. 1998;279(15):1200–5.

7. The importance of pharmacovigilance Safety Monitoring of medicinal products. World Health Organization 2002.

8. Angulo N, Jiménez F, Valcárcel R, Vaca C, Orozco J, López J. Conceptos básicos en farmacovigilancia. Boletín de farmacovigilancia (suplemento) [Internet]. 2006; (3):[1–7 pp.].

9. Angulo N, Jiménez F, Valcárcel R, Vaca C, Orozco J, López J. Conceptos básicos en farmacovigilancia 1. Boletín de farmacovigilancia [Internet]. 2006; (4):[1–10 pp.].

10. Baciu A, Stratton K. The Future of Drug Safety: Promoting and Protecting the Health of the Public. Washington (USA): Committee on the Assessment of the US Drug Safety System; 2006.

11. Aranaz Andrés JM, Aibar Remón C, Vitaller Burillo J. Estudio Nacional sobre los Efectos Adversos ligados a la Hospitalización. ENEAS 2005 Madrid, España: Ministerio de Sanidad y Consumo; 2006. Available from: https://www.msssi.gob.es/organizacion/sns/planCalidadSNS/pdf/excelencia/opsc_sp2.pdf.

12. Institute of Medicine Committee on Quality of Health Care in A. To err is human. In: Kohn LT, Corrigan JM, Donaldson MS, editors. To Err is Human: Building a Safer Health System. Washington (DC): National Academies Press (US); 2000.

13. Organización Mundial de la Salud. Alianza Mundial para la Seguridad del Paciente: La investigación en seguridad del paciente. París (Francia); 2008. [Available from: http://www.who.int/patientsafety/information_centre/documents/ps_research_brochure_es.pdf]

14. Ministerio de salud y protección social. Seguridad del paciente y la atención segura. Bogotá (Colombia); 2014 [Available from: https://www.minsalud.gov.co/sites/rid/Lists/BibliotecaDigital/RIDE/DE/CA/Guia-buenas-practicas-seguridad-paciente.pdf.]

15. Ministerio de salud y protección social. Sistema Único de Acreditación Bogotá (Colombia); 2014 [Available from: https://www.minsalud.gov.co/salud/PServicios/Paginas/sistema-unico-acreditacion-sistemaobligatorio-garantia-calidad.aspx.]

16. Aranaz-Andrés J, Aibar-Remón C, Limón-Ramírez R, Amarilla A, Restrepo FR, Urroz O, et al. Prevalence of adverse events in the hospitals of five Latin American countries: results of the ‘Iberoamerican study of adverse events’ (IBEAS). BMJ Qual Saf. 2011;20(1):1043–51.

17. Velasco-Rico J. Neoliberalismo, salud pública y atención primaria: Las contradicciones en el paradigma de salud para todos. Colombia Med. 1997;28(1):27–33.

18. Menendez EL. Modelo Médico Hegemónico: Reproducción técnica y cultural. Natura Medicatrix. 1998;51(1):17–22.

19. Bailey C, Peddie D, Wickham ME, Badke K, Small SS, Doyle-Waters MM, et al. Adverse drug event reporting systems: a systematic review. Br J Clin Pharmacol. 2016;82(1):17–29.

20. World Health Organization. Marco Conceptual de la Clasificación Internacional para la Seguridad del Paciente. Ginebra (Suiza): Organización Mundial de la Salud; 2009. [Available from: http://www.who.int/patientsafety/implementation/icps/icps_full_report_es.pdf.]

21. World Health Organization. The WHO Programme for International Drug Monitoring Uppsala: World Health Organization; 2017 [Available from: http://www.who.int/medicines/regulation/medicines-safety/about/drug_monitoring_prog/en/.

22. ATC/DDD Index 2014 Norwegian: WHO Collaborating Centre for Drug Statistics Methodology.; 2018 [Available from: https://www.whocc.no/atc_ddd_index/]

23. Ministerio de Sanidad, Política Sociale Igualdad. Incidentes y eventos adversos en medicina intensiva. Seguridad y riesgo en el enfermo crítico. SYREC 2007. Madrid, España: Ministerio de Sanidad, Política Social e Igualdad: 2010 [302]. [Available from: https://www.seguridaddelpaciente.es/resources/documentos/syrec.pdf.]

24. Napal V. M. G, Ferrándiz JR. Dispensación con intervención previa del Farmacéutico: dosis unitarias. 2002 [cited enero 10 de 2018]. In: FARMACIA HOSPITALARIA - TOMO I [Internet]. España: Sociedad Española de Farmacia Hospitalaria, [cited enero 10 de 2018]; [390-414]. Available from: https://www.sefh.es/bibliotecavirtual/fhtomo1/cap2611.pdf.

25. Castro-Espinosa J. Frecuencia y caracterización de los errores de medicación en un servicio de hospitalización de una clínica en Cali, Colombia. Rev colomb cienc quim farm. 2013;42(1):5–18.

26. Machado-Alba JE, Londoño-Builes MJ, Echeverri-Cataño LF, Ochoa-Orozco SA. Adverse drug reactions in Colombian patients, 2007-2013: Analysis of population databases. Biomedica 2016;36(1):59–66.

27. Elliott RA, Camacho El, Campbell F, Jankovic D, Martyn St James M, Kaltenthaler E, et al. Prevalence and economic burden of medication errors in the NHS in England. Inglaterra: Manchester Centre for Health Economics; 2018.

28. Varallo FR, Guimaraes Sde O, Abjaude SA, Mastroianni Pde C. [Causes for the underreporting of adverse drug events by health professionals: a systematic review]. Rev Esc Enferm USP. 2014;48(4):739–47.

29. Pharmacovigilance Risk Assessment Committee. Good practice guide on recording, coding, reporting and assessment of medication errors: European Medicien Agency; 2015 [Available from: http://www.ema.europa.eu/docs/en_GB/document_library/Regulatory_and_procedural_guideline/2015/11/WC500196979.pdf.]

30. Edwards IR. A New Erice Report Considering the Safety of Medicines in the 21st Century. Drug Saf 2017;40(10):845–9.

31. Organización Panamericana de la Salud. Buenas Prácticas de Farmacovigilancia. Washington, D. C. (USA): Organización Panamericana de la Salud; 2011

32. Machado-Alba JE, Giraldo-Giraldo C, Moncada-Escobar JC. Farmacovigilancia activa en pacientes afiliados al sistema general de seguridad social en salud. Rev Salud Pública. 2010;12 (4):580–8

33. Machado-Alba JE, Moncada JC, Moreno-Gutiérrez PA. Medication errors in outpatient care in Colombia, 2005-2013. Biomédica. 2016;36:251–7.

34. de las Salas R, Díaz-Agudelo D, Burgos-Flórez FJ, Vaca C, Serrano-Meriño DV. Adverse drug reactions in hospitalized Colombian children. Colombia Médica: CM. 2016;47(3):142–7.

35. Moscoso-Veloza SM, Ramírez-Cubillos GF, López-Gutiérrez JJ, Gerena-Useche BE. Reacciones Adversas a Medicamentos en el Hospital de Suba de Bogotá. Revista de Salud Pública. 2006;8:209–17.

36. Chaves M. Caracterización de reacciones adversas a medicamentos en adultos mayores de 44 años en Bogotá, D.C., enero a diciembre, 2012. Biomédica. 2015;35:34–42.

37. WHO. Reporting and learning systems for medication errors: the role of pharmacovigilance centres. Switzerland: World Health Organization; 2014.

38. Orozco J. De la farmacovigilancia al monitoreo crítico de los medicamentos: El proceso de registro de medicamentos en Colombia. (Tesis doctorado en salud pública). Universidad Nacional, Bogotá, Colombia; 2012.

39. De-Abajo FJ. El medicamento como solución y como problema para la salud pública. una breve incursión a los objetivos de la farmacoepidemiología. Rev Esp Salud Pública. 2001;75(4):281–4.

40. Laporte JR. Fifty years of pharmacovigilance - Medicines safety and public health. Pharmacoepidemiol Drug Saf. 2016;25(6):725–32.

41. Caster O, Edwards I. Reflections on attribution and decisions in pharmacovigilance. Drug Saf 2010 Oct 1;33(10):805–9. 2010;33(10):805–9.

42. Edwards RI. Causality Assessment in Pharmacovigilance: Still a Challenge. Drug Saf 2017;40(5):365–72

43. Gervas J. El fracaso programado de la farmacovigilancia. A propósito del “escándalo Depakine": hay culpables y no son los médicos. España. 2018 [Available from: http://www.nogracias.eu/2018/03/19/fracaso-programado-la-farmacovigilancia-proposito-del-escandalo-depakine-culpables-no-los-medicos.]

44. Price J. Pharmacovigilance in Crisis: Drug Safety at a Crossroads. Clin Ther. 2018;S0149-2918(18):30088–2.

45. Morin E. Epistemología de la complejidad. Gazeta de Antropología. 2004;20:1–13.

46. Ugalde A, Homedes N. Las reformas neoliberales del sector de la salud: déficit gerencial y alienación del recurso humano en América Latina. Revista Panamericana de Salud Pública. 2005;17:202–9.

47. Illich I. Némesis médica: La expropiación de la salud. Primera edición ed. España: Barral editores; 1975. 218 p.

